# Temporal microbiome road-maps guided by perturbations

**DOI:** 10.1101/049510

**Authors:** Beatriz García-Jiménez, Mark Wilkinson

## Abstract

**Motivation:** There are few tools that allow longitudinal analysis of metagenomic data subjected to distinct perturbations.

**Methods:** This study examines longitudinal metagenomics data modelled as a Markov Decision Process (MDP). Given an external perturbation, the MDP predicts the next microbiome state in a temporal sequence, selected from a finite set of possible microbiome states.

**Results:** We examined three distinct datasets to demonstrate this approach. An MDP created for a vaginal microbiome time series generates a variety of behaviour policies. For example, that moving from a state associated with bacterial vaginosis to a healthier one, requires avoiding perturbations such as lubricant, sex toys, tampons and anal sex. The flexibility of our proposal is verified after applying MDPs to human gut and chick gut microbiomes, taking nutritional intakes, or salmonella and probiotic treatments, respectively, as perturbations. In the latter case, MDPs provided a quantitative explanation for why salmonella vaccine accelerates microbiome maturation in chicks. This novel analytical approach has applications in, for example, medicine where the MDP could suggest the sequence of perturbations (e.g. clinical interventions) to apply to follow the best path from any given starting state, to a desired (healthy) state, avoiding strongly negative states.

## 1 Introduction

High-throughput sequencing allows us to determine the microbial composition of samples more rapidly than typical bacterial culture techniques. It is also able to identify bacteria which cannot be cultured in laboratory conditions. As a result, it is feasible, using metagenomics to temporally monitor the detailed dynamics of a complex bacterial community, frequently, and over short time-intervals. This allows analysis of changes in microbial composition over time, possible interactions between groups of microbes, and/or the influence of external perturbations. Such data could then be used in the construction of models useful for *in silico* prediction of perturbation-outcomes (Fritz *et al.*, 2013).

An example of where such predictions would be useful is in a hospital critical-care setting. Patients suffering from sepsis often die before traditional bacterial cultures can be returned. As such, with little information about the cause of the infection, a wide breadth of clinical interventions are applied to the patient in the hope that one might prove effective. Unfortunately, the resulting disruption-including to the normal microflora of the patient - often leads to serious complicating illnesses such as pneumonia (Boyd *et al.*, 2014). *In silico* models could provide badly-needed guidance on a specific course of interventions - for example, a personalized and contextually-correct sequence of drugs and/or food - that would lead the patient safely back to a healthy state.

There are few large, publicly available longitudinal metagenomics datasets available (Voigt *et al.*, 2015) that could be used to design such models. Most datasets span short periods of time (weeks), though some span several years (Voigt *et al.*, 2015). Most studies are focused on the human gut, but unfortunately, they seldom include longitudinal metadata (i.e. possible perturbations) associated with each sample.

A recent review (Faust *et al.*, 2015), lists a series of observations made regarding temporal changes in the microbiome that are particularly relevant to the work reported here. First, microbial diversity tends to be stable over time in the same environment. Second, microbial communities evolve through stable states that change due to a) external perturbations (e.g. diet), b) direct modifications (e.g. antibiotics, probiotics) or c) transient perturbations (e.g. microbial interactions). Third, subsequent to a perturbation, the community may return to its original state, or may remain in the new (or another) state. Finally, that communities exhibit a priority effect, where the existence of certain strains will prevent specific other taxa from establishing themselves in a community. These observations are encouraging, in that they suggest that it should be possible to build predictive state-change models. Moreover, they reveal that not all state-changes are possible, thus indicating that a desired state-change might require sequential, planned interventions.

State transition diagrams have appeared in prior metagenomics studies (Gajer *et al.*, 2012; Ding and Schloss, 2014). We propose here to add actions to the edges of these diagrams, and to break-up the complete set of transitions into subsets within which different external perturbations could have influence.

Markov Decision Processes (MDPs) have been used to suggest course-of-treatment within clinical decision support systems (Capan *et al.*, 2015; Sonnenberg and Beck, 1993). We decided, therefore, to apply MDP as a novel approach to the analysis of metagenomics data. Although Brotman *et al.* (2014) used continuous-time Markov models to examine a vaginal microbiome dataset (Gajer *et al.*, 2012) (a dataset which we also examine in this manuscript), their approach differs from our chiefly in that they did not correlate actions/perturbations with state-transitions. Apart from applying a different type of Markov model (not an MDP), their objective was also distinct, in that their model measured temporal transitions between infection/non-infection with human papillomavirus (HPV), estimating transition rates between HPV positive and negative states.

Our approach of using MDPs should allow us to stochastically represent changes in microbiomes states, to identify which perturbations change the microbiome state, with what frequency, and how a goal state could be reached and/or how an undesirable state could be avoided. Our intent is to develop and demonstrate a methodology that will identify the sequence of microbiomes (i.e. OTU vectors) that should be traversed to reach a goal microbiome state, starting from any other state, while avoiding undesirable states. Thereby, we will be able to quantitatively predict the nature and sequence of external perturbations required to achieve a goal.

The main contributions of the manuscript are: 1) considering actions or external perturbations that cause transitions between microbiome states; and 2) given an existing state, to determine the optimal sequence of perturbations to apply to reach or preserve a desired microbiome state.

## 2 Methods

### 2.1 Approach overview

Figure 1 outlines our proposed solution. In summary, to determine the optimal external perturbation to employ to achieve a given state-transition, given a specific starting state, we propose to apply a MDP. We represent a microbiome time series as a state transition diagram with actions (i.e. a ‘road-map’), and solve the MDP to determine the optimal strategy of sequential interventions that will lead the microbiome to a goal state.

**Figure 1:**
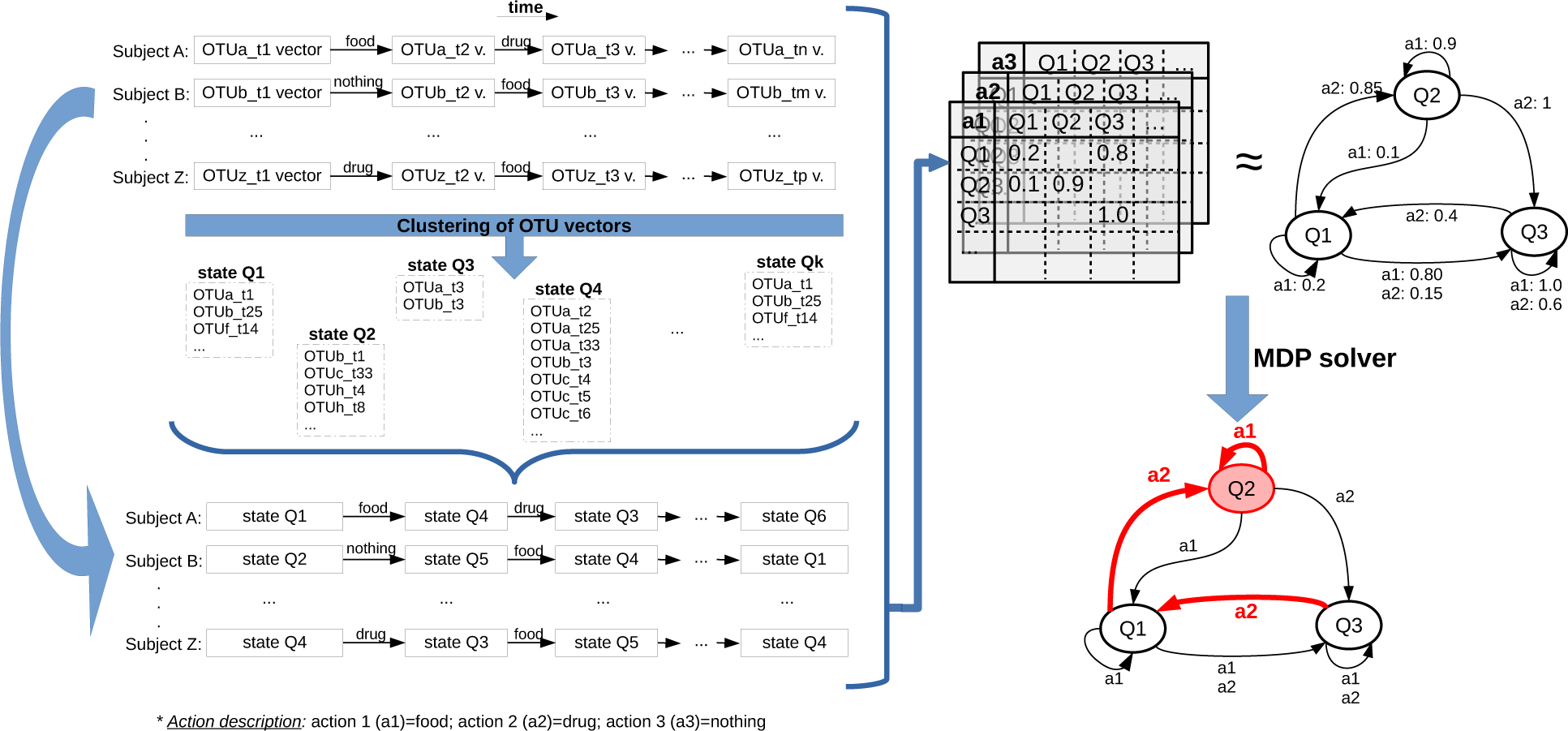
General schema for identifying a temporal sequence of microbiomes depending on perturbations and predicting temporal actions to achieve a defined goal.

Our input is a time series of microbiome data (see Figure 1, top left corner), where each series corresponds to a different subject. Each time point in the series is a microbiome sample, represented by an Operational Taxonomic Unit (OTU) vector, and the number of time points may differ between subjects. Each transition between two time points is labelled with an external perturbation, such as some specific dietary intake, drug, pre/probiotic treatment, etc. That action perturbs the subject microbiome between those two consecutive time points, and therefore is presumed to be the cause of any resulting composition-change.

To create an MDP, we convert the time series of OTU vectors to a time series of microbiome states by means of clustering OTU vectors based on the similarity of their microbial abundance composition. So, MDP states are clusters of OTU vectors, and the transitions, with associated actions, remain the same (see Figure 1, bottom left). Next, from the time series of microbiome states, we obtain a probability transition table for each perturbation/action, and its equivalent transition diagram, labelling each transition with a perturbation and its frequency (see Figure 1, top right). Finally, we apply an MDP solver to identify the path to the goal state through the transition diagram (see Figure 1, bottom right).

### 2.2 Markov Decision Process (MDP)

Markov Processes describe systems with stochastic transitions between discrete-time states. MDP (Bellman, 1957; Howard, 1960; Puterman, 1994) is a type of Markov processes where there are actions to move between states. In a basic Markov Process, the probability of the transitions between states are defined in a bi-dimensional square state-matrix. In an MDP, actions are an added dimension to the matrix. MDP stochasticity allows various transitions to occur from any given state, even given the same action.

An MDP is formally defined as a tuple 〈*S,A,T,R*〉, where:

- *S* is a set of finite states
- *A* is a set of finite actions
- *T:S × A → P(S) = Pr(s*_*t*+1_|*s*_*t*_, *a*), *∀s ∈ S, ∀a ∈ A*
- *R(S)* → *ℜ*

*T* corresponds to the transition probabilities - the probability of going from *s*_*t*_ to *s*_*t*+1_ in the next time slot, given an action *a*. This probability is defined for every triplet of two states and one action. *R* is the reward, and is a real-value depending on the state *s*, representing how “good” that state *s* is. A common alternative definition is *R*: *S* × *A → ℜ* (Puterman, 1994), where the reward is associated to the transition, rather than to the state; and *R(s, a)* is the reward obtained after taking the action *a* from the state *s*. Sometimes, an additional element is also included in the MDP definition: *γ* ∈ [0,1[; called the discount factor, it is a constant typically close to 1.

A solution to an MDP is a policy *(π: S → A)*, i.e., a mapping from states into actions that is used as the basis for decision making. The optimal policy π* for an MDP is the policy that maximizes the expected reward. The optimal policy is usually computed with a dynamic programming algorithm such as value iteration and policy iteration (Bellman, 1957; Howard, 1960).

### 2.3 MDP for temporal metagenomics

Our problem is represented as a generic *MDP=〈S,A,T,R〉* as follows:

- *S* = *{group of microbiome samples}* Goal states ∈ *S*
- *A = {perturbation}*
- *T = {Pr(stateOut | stateIn,action)}, ∀ 〈stateIn, action, stateOut〉 = Frequency (〈stateIn, action, stateOut〉), ∀ stateIn/Out ∈ S, ∀ action ∈ A*, where Σ_*s*_*out*^*∈S*^__ *Frequency (stateIn, action, s*_*out*_) = 1
- 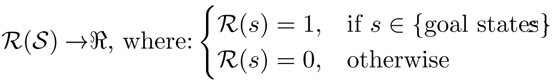

Applying our MDP proposal to find pathways through the resulting ‘road-map’involves: preprocessing; defining the four elements of the MDP (states, actions, transitions and goal states for rewards); and solving the MDP. These steps are described in the following subsections.

#### 2.3.1 Step 1: Pre-processing microbiome data

Metagenomics pre-processing steps are necessary, but not standardized, being highly variable from laboratory-to-laboratory. Pre-processing is dependent on the specific methodologies/technologies for sample preparation and data generation, and cannot be prescribed in a generic manner for all datasets. This is particularly relevant for this study, since we utilize datasets generated by three other laboratories, each with its own intrinsic preprocessing requirements.

The pre-processing steps described below were applied in our first two datasets (human and chick gut microbiome, see section 2.4.1 and 2.4.2), but not the third (vaginal microbiome, see section 2.4.3) because the prepared data was available online.

Our pre-processing followed the methodology of David *et al.* (2014). It includes removing OTUs do not present in any samples (from other body sites, other donor, etc.); removing samples due to suspicion of experimental noise or contamination (as defined per experimental procedure); removing samples with low read-counts (less than 10,000); and normalizing the OTUs (see below). Additionally, it is sometimes useful to aggregate taxa at higher taxonomic level than species, for example, aggregation at the genus level as was done by Larsen and Dai (2015) for the David *et al.* (2014) dataset.

Our pre-processing follows the steps described in the section ‘Sample quality control’of David *et al.* (2014). It includes removing OTUs whose sum is 0 (from other body sites, other donor, etc.); removing samples due to suspicion of experimental noise or contamination (as defined per experimental procedure; for example in David *et al.* (2014), but not in Ballou *et al.* (2016)); removing samples with low read-counts (less than 10,000); and normalizing the OTUs (see below). As additional pre-processing, it is sometimes useful to aggregate taxa at higher taxonomic level than species, for example, aggregation at the genus level as was done by Larsen and Dai (2015) for the David *et al.* (2014) dataset.

One of the most influential pre-processing steps is normalization of the OTU counts. We have selected *David et.al*.’*s* normalization (David, 2014), because it includes a log transformation which is recommended to preserve the relative microbe relations, taking into account the compositional nature of the data (Aitchison, 1982). We use the python code published by the authors (David, 2014) but we add a pseudo-count with “a value smaller than the minimum abundance value prior to transformation”(Costea *et al.*, 2014). In particular, the selected pseudo-count is one order of magnitude (base 10) less than the minimum abundance value, as recommended by Costea *et al.* (2014). For example, for a minimum abundance value of 1.623e-06, the pseudo-count will be 1e-07.

#### 2.3.2 Step 2: Defining MDP states

The set of possible microbiome states clearly must be simplified, since the total space of the OTU vector is so large as to not be computationally tractable for MDP solutions. However, oversimplification results in small numbers of states, which are insufficiently granular to measure anything “biologically meaningful”. Therefore, we must consider this when defining an approximation to group similar OTU vectors.

*How to deterministically convert an OTU vector into a MDP state:* Our definition of MDP states involves a standard clustering-of-samples. Historically, clustering in metagenomics applies different approaches, in terms of distance measure, algorithm and number of clusters. As distance measure, the Jensen-Shanon Distance (JSD) (or its root squared, as in Arumugam *et al.* (2011a)) is the most frequently used (Gajer *et al.*, 2012; Ding and Schloss, 2014); although the cophenetic or the Euclidean distance are sometimes applied. Several clustering algorithms have been used to group metagenomics samples, such as PAM, Agnes, Hclust, or Dirichlet Multinomial Mixture; with different linkage options. For determination of the number of clusters, diverse assesment criteria have been used in the literature: the average Silhouette width (SI), Calinski-Harabasz (CH) index, Laplace approximation, etc.

For example, a clustering of samples was applied to define enterotypes (Arumugam *et al.*, 2011a). According their tutorial (Arumugam *et al.*, 2011b), they computed the distance as the root square of the JSD, with the PAM algorithm and selecting the number of clusters with the CH index combined with a SI assessment. On the other hand, Gajer *et al.* (2012) applied hierarchical clustering with JSD, and SI assessment. Of these, the latter is the most similar to our selected approach.

Our selected procedure consists of applying Agnes (Kaufman and Rousseeuw, 1990), a hierarchical clustering algorithm, taking the JSD beta diversity metric as the distance measure between samples (with its weighted average linkage to measure the distance between clusters), and SI as the criterion for selecting the number of clusters. Clustering and the distances between OTU vectors were computed with the R implementation of Agnes (Maechler *et al.*, 2015), and the distance function from the R phyloseq package (McMurdie and Holmes, 2013).

Our choice of clustering parameters was guided by the desire to identify several (>2) well-populated microbiome states both within and between individuals, which is a goal disparate from the more common research-aim which considers the stable-state microbiome as a single entity (as with the enterotype studies). As such, the ‘default’ clustering parameters that appear in most of the published approaches do not mirror our problem requirements. We are interested in more than two, relatively well-populated clusters, which provides sufficient heterogeneity in MDP states that we can build a richer, wider, more granular, and (we believe) potentially more biologically-interesting MDP model.

To tune the clustering parameters to our MDP modelling requirements, we attempted two alternative procedures. The first was taken from Gajer *et al.* (2012), that searches for the *k* number of clusters with the best SI being *k* limited to a range of 2 to 10. When this procedure results in a single cluster (MDP state) with a size equal to or greater than 5 samples, we attempt to get more well-populated MDP states using the second approach. This second approach involved selecting the nearest *k* with an SI greater than 0.25 (the minimum threshold for ‘sensible’clusters, according to Rousseeuw (1987)).

Subsequent to either approach, we added an additional step that avoids clusters/states with very few samples (less than 5 samples), redistributing the instances of these very small clusters to the nearest cluster according to the original Agnes results. If that nearest cluster reported by the Agnes R object is also a removed cluster, due to its small size, the sample is moved to the first large cluster of the nearest sample, according to the previously computed JSDs, repeating this procedure until a neighbour sample belonging to a large preserved cluster is found.

#### 2.3.3 Step 3: Defining MDP actions

This is the primary contribution of this manuscript. Including perturbation actions in microbiome analysis has not been reported in any of the (relatively few) prior studies that discuss microbiome state-transitions. The definition of what constitutes an ‘action’depends on the particular experimental question, and so we define it independently in each of our analysed datasets below 2.4. Briefly, actions can be nominal, for example, salmonella vaccine and probiotic in chicks (see section 2.4.2); numerical after discretizing, such as nutritional intake in human gut (see section 2.4.1); or binary, for instance, sexual practices (see section 2.4.3). Since the set of actions in an MDP must be finite, we must discretize any quantitative and continuous values in perturbations. Finally, for simultaneous or concurrent actions, we represent and solve a different MDP for each. For example, an MDP for the *{low, medium, high}* fiber intake, another MDP for the *{low, medium, high}* fat intake, etc.

#### 2.3.4 Step 4: Defining MDP transitions

Transition probabilities *T* are built on a set of triplets taken from the microbiome temporal sequences, by looking for the preceding and subsequent states of every external perturbation. The microbiome sequence of each subject is split into as many triplets as there are perturbations, where the input (respectively, output) state of the triplet is the microbiome sample before (resp., after) the perturbation occurs. So, triplets *〈input-state, action, output-state〉* come from individual steps in the microbiome time series. Finally, the frequency of this triplet is computed.

#### 2.3.5 Step 5: Defining MDP goal states

Our reward modelling depends only on states. *R* is based on either desirable ‘goal’state(s) or on ‘undesirable’state(s). In the latter case, *goal states=S-{undesirable states}*. However, a more specific reward schema could be defined, depending also on the action per state, if the microbiome data includes sufficiently detailed meta-data regarding interventions.

#### 2.3.6 Step 6: Solving the microbiome MDP

Here we apply a search algorithm to identify a policy. We use the *MDPtoolbox* (Chads *et al*, 2014) R package (version 4.0.2) to solve our MDPs. In our scenario, the time horizon over which decisions need to be made by the MDP is indefinite horizon, because we have some terminal states that correspond to our goal (e.g. a healthy state, or the state/s with the highest microbial diversity). *MDPtoolbox* allows rewards depending on transitions. Among the available algorithms and optimization criteria (Chads *et al.*, 2014; Puterman, 1994), we have selected the policy iteration (Howard, 1960), because it requires fewer iterations; however any algorithm that converges could provide us with a valid policy. We selected a discount factor 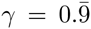. There could also be more than one optimal policy. For example, given a state *Q1*, when the application of two different actions *a1* and *a2* from *Qi* results in exactly the same reward, both strategies *a1* and *a2* are equally good for *Qi*.

### 2.4 Datasets

Few public longitudinal microbiome datasets include sufficiently frequent sampling and associated meta-data to allow perturbation studies. Nevertheless, we found three distinct datasets which can be modelled as MDPs according to our proposed methodology, to determine the perturbation influence on microbiome dynamics.

#### 2.4.1 Human gut microbiome

We analyzed a dataset from David *et al.* (2014), the longest longitudinal study, as far as we know, about the gut microbiome in healthy subjects, which has also been used in other posterior studies (Larsen and Dai, 2015). It consists of near-daily saliva and stool sampling of two subjects, donor A and donor B, along a whole year (i.e. our temporal microbiome sequences). It includes meta-data taking note of their dietary intake, which for us becomes the external perturbation for the gut microbiome. Thus, this datasets fulfils the basic data-requirements of our approach. The input OTU table with absolute abundances was kindly shared by the authors in a personal communication. The other required input for our MDP model was the nutritional meta-data, taken from the additional file 18 in the original publication (David *et al.*, 2014). Additional information about the study indicated that there were short periods of dysbiosis: in donor A because of travel abroad and in donor B due to a gut illness, with a posterior microbiome recovery.

The MDP model for the human gut domain is *MDP*_*hg*_ = 〈*S,A,T,R*〉, where:

- *S* = *{clus1, clus7}*
- *A*_*i*_ = *{low, medium, high} x {calcium, calorie, carbohydrates, cholesterol, fat, fiber, protein, saturated fat, sodium, sugar}*
- *T*_*i*_: Transition probability tables, with 132 transitions each, available online.
- *R:* Goal state = *{clus7}, if(goal state) then R=1; else R=0*

With all the 493 gut samples with 4746 taxa, including both subjects, we obtained 7 clusters (see supplementary Figure S1(a)) using the clustering procedure described in section 2.3.2, which requires that we reach, at least, an SI of 0.25. Unfortunately, after removing samples without available perturbations (i.e., those that lack metadata about the nutritional intake of the day preceding that sample) only 136 samples remained. These samples populated only 2 clusters (see Figure S1(b)), all of them belonging to donor A. As such, we could determine influences only for this donor. We have 2 MDP states, where most samples belong to the same state (86.76% in cluster 1). 15 samples from cluster 7 correspond to days 15, 19, 26, 41, 42, 59, 136, 163, 192, 197, 198, 199, 200, 201 and 202, 60% of them falling in the period after dysbiosis (travel from day 71 to day 122). As such, the distinct clusters could be due largely to this period of travel, and have little greater biological significance.

Regarding the perturbations *A*_*i*_, since the nutritional intakes of different elements are simultaneous, we defined an MDP for each of the 10 nutritional elements available in the meta-data. In this domain, we decided to discretize in three different bins, split by 0.33 and 0.66 thresholds, resulting in 3 different actions: low, medium, high.

To define the reward *R*, we need to know the goal state/s. Unfortunately, after the filtering steps above, the days which the original study describes with diseases or microbiome dysbiosis are not included in the remaining samples. Thus, for the purposes of our study, we define as goal state the cluster 7, that one within more samples after dysbiosis than before, since the ‘goal’and/or ‘avoidance’states can be defined arbitrarily (with respect to creating the road-map).

#### 2.4.2 Chick gut microbiome

This dataset was generated by the recent study of Ballou *et al.* (2016). It analyses the response of different treatments (salmonella vaccine and/or probiotics) in the chick gut, during their first month of life. It includes 119 samples with 1583 taxa. The samples include six time points (days 0, 1, 3, 7, 14 and 28), with approximately 4 or 5 subjects per 4 treatments.

The MDP model for the chick gut is *MDP*_*cg*_=〈*S,A,T,R*〉, where:

- *S = {Q1, Q2}*
- *A = {cc, cp, sc, sp}*, where c=control (not salmonella and/or not probiotics), p=probiotics, s=salmonella
- *T*: Transition probability tables, with 94 transitions, available online.
- *R:* Goal states= *{Q1}, if(goal state) then R=1; else R=0*

For states *S*, we were able to follow Gajer *et al.* (2012)’s criteria to determine the clusters, resulting in 2 final MDP states.

The actions *A* are 4 different treatments, corresponding to the 4 combinations of 2 external perturbations: Salmonella vaccine and/or probiotic supplement (0.1% Primalac). The action is the same for the same subject in each time series. Salmonella vaccine was given at the beginning, before the day 0 sampling, to chicks in the group ‘sc’and ‘sp’. Probiotic was mixed with the food, on every day of the experiment, only to the groups ‘cp’and ‘sp’.

The reward *R* was determined, first, by asking the study authors if they had defined “healthy” versus “non-healthy” chicks. They indicated that this was not a factor in their experiment (that all chicks were healthy), and as such, we defined the goal states using the default criterion: the highest mean alpha diversity, before normalizing, according to the Shannon metric, because higher diversity is generally correlated with healthier states.

The data was downloaded from the Qiita repository *(http://qiita.ucsd.edu)*, with study no.10291. The OTU table is in biom format (McDonald *et al.*, 2012), and the mapping data is contained in a text file.

#### 2.4.3 Vaginal microbiome

The MDP model for vaginal microflora is *MDP_v_ = 〈S,A,T*,R*〉*, where:

- *S = {I, II, III, IV-A, IV-B}*
- *A*_*i*_ = *{ Yes, No}* x {anal sex, digital penetration, douching, lubricant, oral sex, sex toy, tampon, vaginal intercourse}
- *T*_*i*_: Transition probability tables, with 905 transitions each, available online.
- *R:* Non-desired state= *{IV-B}, if(non-desired state) then R=0; else R=1*

This dataset come from Gajer *et al.* (2012). The OTU table and the clusters come from supplementary table S2 (Gajer *et al.*, 2012), which consists of 937 samples and 330 OTUs, corresponding to 32 women collecting samples twice per week for 16 weeks. The data counts are already pre-processed, and normalized to a sum of 100 per sample, being relative abundances.

State type ‘communities’were defined by the original study authors, and they also label our clusters/MDP states, *S* (available in the supplementary Table S2 (Gajer *et al*, 2012)). Gajer *et al* obtained the clusters by hierarchical clustering, with Jensen-Shannon distance, with Ward linkage, cutting the dendogram with a *k* between 2 and 10, with the maximum silhouette inside this range. The maximum silhouette was at *k*=5, and thus they obtained 5 states.

We computed the JSD with this matrix to check the SI value of these clusters, and to plot some graphs to determine where the samples with bacterial vaginosis are located, this being defined as the ‘avoidance’state.

The actions set *A* was composed by the available meta-data related to the hygienic and sex activities that could perturb the vaginal microbiome. These actions were collected by a curated visual inspection of the individual profiles of the dynamics of vaginal bacterial communities, from the 32 D-panels (one profile per woman) of supplementary material in Figure S5 (Gajer *et al*, 2012) available on-line. We associated the external perturbation to the next sample taken, or the same day if it coincides, and this is then considered the ‘action’between the two samples.

This MDP analysis differs from the previous two in that the reward *R* is defined differently, being based on a non-desired state rather than a goal state. In this scenario, we want to avoid Bacterial Vaginosis, so this is the state with the lowest reward. The Nugent score, from microscopic bacterial observation, is typically used as the diagnostic for bacterial vaginosis (when it takes a high value) (Nugent *et al.*, 1991). The quantitative Nugent score has previously been discretized into *low (0-3), int(ermediate) (4-6), high (7-10)*. Therefore, these values were used to assign the rewards in our MDP. The non-desired state - the state to avoid-is IV-B, which concentrates most of the high and intermediate Nugent categories (see supplementary Figure S2), which indicates the greatest risk for the Bacterial Vaginosis disease. The remaining states are considered equivalent (non-diseased), and assigned a reward of 1.

## 3 Results

### 3.1 Human gut microbiome

Despite the sparsity of meta-data about external perturbations, as proof of concept, we could solve the *MDP*_*hg*_ defined in the section 2.4.1, obtaining a transition diagram and a policy for each different perturbation (fat, protein, fiber, and so on), that show how the different nutritional intakes could have influence on the human gut microbiome dynamics. According to Hekstra and Leibler (2012), temporal relations can be inferred from: a) many replicates from the same time point, or b) one replicate at many time points. So, even with only one subject with many sequential observations, as remained after the filtering of David *et al.* (2014), we could nevertheless retrieve biologically informative patterns.

We notice a high degree of stability in the human gut microbiome for donor A. We could consider that the most populated state (cluster 1) includes the days of normal diet, and cluster 7 the days after the change in diet and recovery resulting from travel to a developing Southeast Asian country. Cluster 7 is, however, very similar to cluster 1, despite constituting more samples from the recovery period than the pre-travel period. By modelling the problem with an MDP, where nutritional intakes are the perturbations or actions to choose in order to reach microbiome state 7 (the goal state), we could interpret the optimal policy suggested by the MDP solver in state 1 (see supplementary Table S2) as being equivalent to the nutritional qualities of the foods eaten while abroad.

Though there are several noteworthy observations from this analysis, we will comment only on one exemplar - that is, with fiber intake as perturbation (see Figure 2(a)). The objective is to reach our selected goal state, *clus7*, with a majority of samples from the recovery process. Starting from *clus1*, the MDP policy indicates that the optimal action (among low, medium or high fiber intake) is to ingest significant amounts of fiber (greater than 21.43gr/day, representing the upper 33% of the range), and it would be successful 14% probability (86% probability of remaining in *clus1*) as the diagram in Figure 2(a) shows. However, if the microbiome is already in the goal state, *clus7*, the optimal action suggested by the MDP policy to maintain that state is to consume a low quantity of fiber, which preserves the goal state with 67% probability. Conversely, with a high quantity of fiber intake, there is an equal likelihood of remaining in *clus7* versus moving to *clus1*.

**Figure 2:**
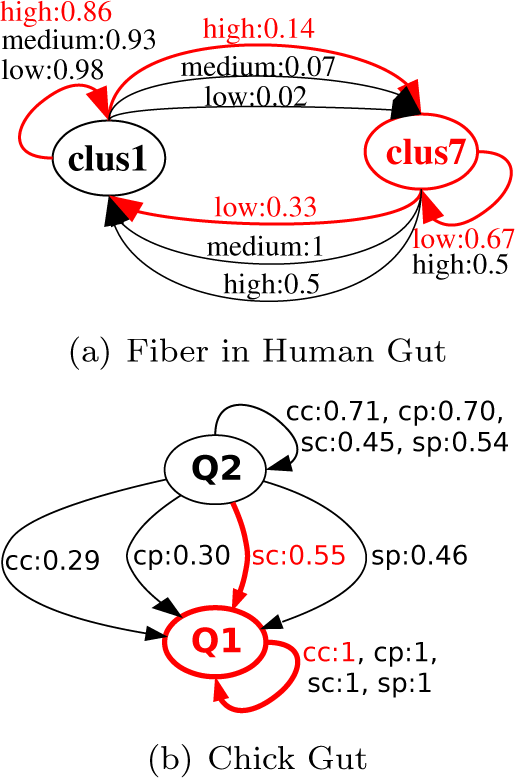
MDP diagrams and solutions to Human and Chick gut microbiomes. Red arrows represent the MDP solution, i.e. the policy returned by the MDP solver. The goal state is that highlighted in red: (a) Fiber-intake as perturbation in human gut microbiome ‘road-map’; (b) The chick gut microbiome ‘road-map’.

### 3.2 Chick gut microbiome

The manuscript describing the Ballou *et al.* (2016) dataset indicates that there are no differences in the health of the chicks at the end of the study. In this section, we examine the possibility of confirming this by discovering transitions between chick microbiome states.

First, we observed that the clusters/states primarily mirror the chicks’ age (i.e. sampling day): there is one state (called *Q2*) with samples from chicks in their first days of life (day 0, 1, 3 and some chicks from day 7), and another MDP state (*Q1*) with microbiomes of all the chicks aged 2 or more weeks (day 14 and 28) and some samples from chicks aged only 1 week (sampling of day 7). This split according to age is in agreement with the Ballou *et al.* analysis (Ballou *et al.*, 2016), where samples do not differ by any other criteria, such as salmonella vaccine administered, or diet including/excluding probiotics. Both states include samples from the four different treatments. Consequently, both states have a similar number of samples *(Q1* with almost the 45% and *Q2* with the remainder 55%), although *Q1* is much more diverse than *Q2* with respect to microbial composition (mean alpha diversity: 3.34 *(Q1)* ≫ 1.48 *(Q2))*.

When we analyse the sample distribution after clustering and divided by treatment, we identified a pattern in the state of the samples from day 7. All chicks without any treatments *(cc)* or with only probiotics *(cp)* after their first week (day 7) are grouped with newborn/baby chicks (state *Q2)*. However, chicks of the same age but only treated with salmonella vaccine and no probiotics *(sc)* are all in state *Q1*, more similar to the microbiome of adult chicks.

This pattern is also identified by our MDP solution, which reveals the shortest path from the current state to the goal state (see Figure 2(b)). On the one hand, *sc* is the optimal perturbation plan to arrive to the goal state *Q1* from *Q2*. On the other hand, the probabilities in the transition diagram confirm that with *cc* or *cp* there is more than a 70% likelihood of remaining in an ‘immature’microbiome state. Therefore, we could con-clude that an adult microbiome, with more diversity, is reached earlier with the salmonella vaccine treatment without probiotics.

Thus, MDP models can reveal interesting, biologically-relevant, and quantitative patterns of microbiome evolution under varying environmental conditions. Moreover, setting diversity as our goal brings our Chick MDP model into agreement with child microbiome evolution studies (Dominguez-Bello *et al.*, 2011; Oakley *et al.*, 2014), which begin with an empty or very low-diversity microbiome (such as our *Q2* state with the youngest chicks), while achieving maximum diversity in adulthood (reached in one or two weeks for chicks).

### 3.3 Vaginal microbiome

This dataset provides more diversely annotated observations than the previous two gut datasets, and we are therefore able to apply our MDP to interpret the rapid fluctuations in the vaginal microbiome. In addition, this scenario differs from the previous two in that most states are goal states; the objective is to avoid arriving at the disease state. For this experiment, sample clusters were adopted exactly as published in the original study (Gajer *et al.*, 2012), and we assigned each MDP state the same label (I, II, III, IV-A, IV-B) as the author-defined community state type for that cluster.

As a general overview, all states, except to IV-A, have more than 85% likelihood of remaining in the same state, regardless of action (see Table S1 with global relative transition probabilities). In addition, there are almost as many output links as input links per state (comparing sums of rows and columns in absolute abundance table).

“Some of the taxa in communities of state type IV-B have previously been shown to be associated with bacterial vaginosis”(Gajer *et al.*, 2012). Therefore, we considered state IV-B as the ‘diseased’, non-desired state, and the MDP was designed to avoid reaching it, or to move from it to a healthy state (state I to IV-A). Therefore, the successful path is the action or series of actions that minimize the risk of reaching state IV-B; effectively, not selecting the action that has the highest transition-frequency that arrives at the diseased state.

Table 3.3 represents a combined behaviour policy to preserve the health in the vaginal microbiome, avoiding bacterial vaginosis. Each column in Table 3.3 represents the policy about how to act, depending on the vaginal microbiome state type that each woman has. Table 3.3 shows that the recommended perturbations are entirely dependent on the current microbiome state, as there are no perturbations for which the recommended value is the same for all the states. However, with a microbiome in state III or IV-A any perturbation is a valid MDP policy to preserve health.

**Table 1:** Combined recommendations to avoid bacterial vaginosis. Rows represent external perturbations that could influence the vaginal microbiome. Columns represent the microbiome states. In each cell, the policy (yes/no) derived from all Markov systems, one per external perturbation.

| Perturbation\State | I | II | III | IV-A | IV-B |
| --- | --- | --- | --- | --- | --- |
| **Anal sex** | No | No | Yes | Yes | No |
| **Digital penetration** | Yes | No | Yes | Yes | Yes |
| **Douching** | Yes | No | Yes | Yes | Yes |
| **Lubricant** | Yes | No | Yes | Yes | No |
| **Oral sex** | No | Yes | Yes | Yes | Yes |
| **Sex toy** | Yes | No | Yes | Yes | No |
| **Tampon** | Yes | Yes | Yes | Yes | No |
| **Vaginal intercourse** | Yes | No | Yes | Yes | Yes |

Figure 3 shows that the most common model-behaviour is to maintain the same state, regardless of perturbation, and state-to-state transitions are less common (thiner ribbons). For example, when perturbations take a value ‘No’(bottom row), the profile of transitions is almost the same, regardless which perturbation was avoided; and very similar when all transitions are represented together (see Figure S4 from Gajer *et al.* (2012)). However, state-to-state transitions differ depending on the perturbation applied (see top row in Figure 3), and also differ from the corresponding pair with perturbation=‘No’(pairs in the same column). Take, for example, the data for anal sex and lubricant-use. The near-absence of transitions (very thin lines) going out state IV-B (diseased) when either anal sex or lubricant take a value ‘Yes’, shows the difficulty to exit the state associated with bacterial vaginosis; alternately, we note the high probability of transition from state IV-A to state I or state III when digital penetration takes value ‘Yes’.

**Figure 3:**
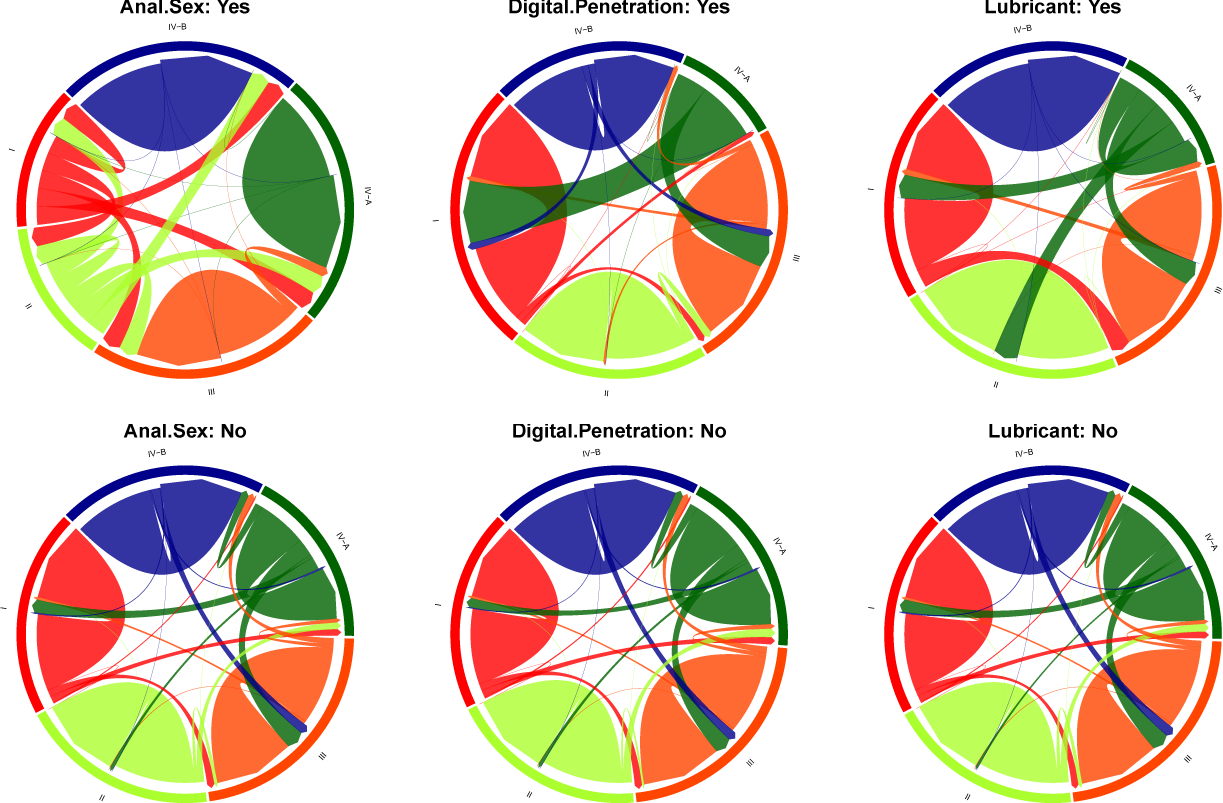
Graph representation of vaginal microbiome state transitions, sorted by external perturbation. The ribbon-arrow points out the directionality of the transitions, and ribbon width represents the frequency of transition. Top row circles show the transition probabilities when the perturbation is applied, and bottom row when it is not. The colors representing each state are I:red, II:light green, III:orange, IV-A:dark green and IV-B:blue. This selection of visualization is adopted from Gajer *et al.* (2012).

Brotman *et al.* (2010) has identified several ‘risk factors’ for bacterial vaginosis, including the use of a lubricant and rectal sex. Our temporal metagenomics MDP approach could perhaps explain this result. When the perturbation is Anal Sex, the most common policy is I: No, II: No, III:Yes, IV-A:Yes, IV-B:No (see diagram in Figure S3). Starting from state I and II this policy is clear because there are no examples of ‘Yes’ transitions (meaning probability=0 (*i.e. Pr=0*). From state III, the probabilities are yes:0.9 and no:0.86. This highly counter-intuitive result appears to suggest that it is (marginally) better to engage in anal sex in order to preserve this healthy state. Such counter-intuitive results arise in part due to lack of sufficient examples in the dataset, and also because this analysis looked at each action in isolation, thus we do not know what other actions were being taken simultaneously. A similarly counter-intuitive rule arises from state IV-A because action=Yes keeps the system in the same state, while with action=no there is a slightly higher possibility to move to the disease state. Finally, from the disease state IV-B, with action=yes (*Pr=1*), the system will persist in the same (diseased) state indefinitely, so the last option is action=no, in order to have some possibility to move to another (healthy) state. These results mirror the results from the original paper -*Pr*(IV-B,yes,IV-B)=1 indicates that persistent anal sex is not recommended for recovery from bacterial vaginosis.

Biologically, it may also be possible to interpret our temporal metagenomics MDP results as a possible prediction of perturbation inducing patterns of interaction between specific microbes present in two states. For example, when the vaginal microbiome is perturbed with the use of lubricant, the MDP shows a transition between state I and state III happens 20% of the time, versus the only ~7% of the time that this happens by default (i.e. with no lubricant or globally with any combination of actions). Figure 3 shows large differences in the frequency of transition from state I (red) to state III (orange) (see the width of the red ribbon): the frequency is high for the policies ‘lubricant:yes’ and ‘anal sex:yes’, but low in the bottom row (‘anal sex:no’, ‘digital penetration:no’ and ‘lubricant:no’) and also with ‘digital penetration:yes’. Thus, comparing the abundance of microbes in the two states I and III involved in the interaction, it could mean that lubricants and anal sex might facilitate the increase of the predominant bacteria in state III to the detriment of the predominant bacteria in state I.

Overall, from our MDP analyses, we provide a quantitative explanation for why anal sex and lubricant often lead to bacterial vaginosis, and/or maintain it. In fact, when they are used, there are a 100% probability of persisting in that disease state. We also note that the use of a sex toy results in a 100% probability of maintaining the disease state. Other perturbations also show a high degree of maintenance of this disease state, suggesting that this state is clearly difficult to escape from without specific medical intervention.

## 4 Discussion

The main contributions of this manuscript are: the inclusion of actions in the state transition diagrams and the prediction of the sequence of external perturbations (i.e. intakes or actions) to preserve or to reach a healthy or desired microbiome state.

Our proposed MDP approach has several positive features. It can be applied to very different datasets (human gut, chick gut, vaginal microbiota), with different qualitative or quantitative external perturbations. In addition, if we interpret each transition between MDP states as a set of interaction patterns, as was shown in the section 3.3, this new approach also could contribute an additional objective of identifying interaction patterns between two OTU matrices.

MDPs allow different representations/interpretations, depending on experimental question, even given the same dataset. For example, you can modify the MDP to optimize for microbial diversity, or you can modify the MDP to optimize for recovery from a vaccination, depending on your requirements. In the chick dataset, an alternative study design to that described above would be to consider salmonella vaccine as criteria to define the states (with/without vaccine), given that it is administered only at the beginning. Similarly, in the vaginal flora microbiome, an alternative representation could model the reward *R* as being associated to the transition instead of the state, using the Nugent category available for each sample. Some samples with high Nugent category (i.e. the sick ones) do not belong to the state IV-B, and numerous samples with low or intermediate Nugent category, are grouped in this denominated non-healthy state IV-B (see supplementary Figure S2).

There is, perhaps, an oversimplification of our study design in the case where the tampon is the external perturbation. This case takes into account the use of tampon just during the menses. Given that menstrual cycles are the primary driver of vaginal microbial dynamics in women (David *et al.*, 2014) referring to in Gajer *et al.* (2012), a better MDP design for this scenario might be to include other variables (such as menses/no menses) in the state definition. For example, duplicating the MDP states for each of the current states (I to IV-B), representing state X with menses and state X no-menses. Doing this, however, would make the problem larger and more data would be required to cover all cases with sufficient frequency.

There are other approaches, that we did not pursue, to describe the MDP states. One could define as many states as combinations of the possible values of a selected subset of OTUs. For example, if we select 3 OTUs, with 2 possibles values (presence/absence), we will have 3^2^=9 states, or with 10 OTUs with 2 values, 10^2^=100 states. Alternatively, the selection of OTUs could be determined by a variety of criteria: a) by highest frequency; b)by goal states: OTUs expected to be in the discriminant states; c) based on literature, e.g., OTUs known as the most relevant in a modelled disease; d) based on the most relevant/characteristic OTU, e.g., OTUs which are output from a feature selection algorithm from Machine Learning; or e) based on microbial associations, e.g., OTUs with microbial associations to each-other.

Selection of appropriate states requires careful consideration, in particular because the Markov property requires that the next state depends only on the current state and the current action, regardless of previous states and actions. In our temporal microbiome scenario, this might not be completely true (since interventions, such as food or drug intake, can have delayed effects that last over many observational cycles). Nevertheless, on this study we consider this simplification to be quite minimal, and that in general the predictive power gained by the MDP principles is worth this potential limit or noise in its sensitivity.

One drawback of our approach is there is not a deterministic state/cluster definition that provides a reliable split of metagenomics samples into multiple states, as would be desirable for our MDP. The main reasons for this could be the scarcity or lack of variability in the data, as was the case with gut microbiota composition of which two of our three datasets are comprised. With more diversity in the data, as seen in the vaginal samples, this becomes less problematic and several states of interest can be identified. In quantitative terms, we take the Silhouette coefficient (SI) as assessment measure of clustering results, where the minimum threshold of acceptable clusters is 0.25. Thus, the human gut dataset has a SI value very low, even more-so after regrouping isolated samples in bigClusters (mean=0.33 with 7 states, and mean=0.22 limited to 2 states with samples with available action data). The SI of chick gut data is better (mean=0.42), and the SI of the vaginal microbiota data is good (mean=0.63).

We also cannot be sure that the different external perturbations that compose a macro-policy need to be applied all together to preserve the health, nor can we can assert that the individual application of only some of the suggested external perturbations be enough to preserve the health. We obtain a policy independently for every perturbations, however in the real data multiple perturbations could be (and were) applied concurrently. Therefore, the predicted policy by the MDP of a particular perturbation *A*, could be affected by additional perturbations *B, C* and/or *D*, and therefore might not have the expected result. To take into-account the effect of all the possible perturbations simultaneously, we would need to build an MDP where the actions represent combined perturbations, and we would need dramatically more data to cover all such combinations.

As with most population-based studies, the quality of the transition diagrams are highly dependent on the quality and quantity of the available data. Some pair (transition,action) could be built based on only a few, or certain important (transition,action) could be missed entirely. This lack of sufficient examples leads, for example, to the highly counter-intuitive plans recommended for anal sex in the vaginal flora study.

Despite these drawbacks, there is a clear and pressing need to predict the effects of interventions on microbiomes, both within medicine and for environmental engineering. This paper provides, to our knowledge, the first example of a MDP being used to explain (and in principle, predict) how a microbial community will respond to any given intervention. We believe this provides the basis for a wide range of directed research, in particular with respect to microbiome-engineering for health in the medical domain.

## 5 Conclusion and Further Work

The proposed method builds a model that suggests external perturbations that should be applied to a given microbiome state, resulting in its navigation through a subset of healthy or acceptable states, avoiding disease or other non-desirable microbiome states. This manuscript confirms that, given sufficient data, this method can be applied to a diversity of temporal metagenomics datasets, such as the three distinct domains (human gut, chick gut and vaginal microbiota) whose solution in terms of microbiome dynamic was found using the proposed MDP strategy.

Integrating the external perturbations together into a single transition diagram is a clear and obvious next-step. In that case, a transition between states could be labelled with more than one perturbation, which more closely represents ‘reality’; for example, most patients are both taking drugs, and eating foods that include fats, proteins, and fibers, so considering them independently will surely miss important effects. To achieve this requires both a dramatic increase in the amount of data collected, which should include subjects spanning all combinations of treatments or behaviours, and a strict adherence to a meta-data collection policy by the researchers.

When the quantity of publicly-available, richly annotated microbiome time series increase, many different future works will be enabled. For example, expanding into plants and soil microbiomes, where temporal data is very limited despite a myriad of interesting external perturbations such as natural and artificial fertilization, crop-rotation, pollution, water toxicity, and so on, that could affect the microbiome composition of soils, root and leaves. With increasing data, new types of actions could be considered, with the most exciting being to consider microbial interactions themselves as transient perturbations, as Faust *et al.* (2015) suggested. Further to this, we might take into account more than one microbiome source, such as different body cavities, where the distinct populations affect each other indirectly, but otherwise behave (we assume) relatively independently.

Finally, one additional future work would be to generate new simulated datasets using the MDP transition model, generating random numbers and applying in each case the corresponding action from each state. If we could computationally synthesize ‘realistic’microbiome time series in a massive way, and parametrically change the properties of the dataset in terms of, for example, the number of samples, taxa, time slots, frequency in the time slots, etc., it would be possible to generate microbiome temporal datasets that reflected a scientifically-hypothesized ‘reality’. This would then allow us to determine, for example, the minimum cohort-size for a proposed study to investigate that hypothesis, where the researcher has a general idea of the number of parameters, and the strength of their effects. In an age where many studies prove to be non-reproducible largely as a result of insufficient data, determining cohort size a priori will become an increasingly important undertaking, particularly for complex studies such as the ones proposed here.

## Acknowledgements

Thanks to the authors of David *et al.* (2014) and Ballou *et al.* (2016) for kindly answering our requests about their datasets. We also thank Dr. Keith Walley for his stimulating explanations of the clinical need for these kinds of algorithms.

## Availability

Referred supplementary figures and tables (i.e.*Sx*) are at BioRxiv data supplements. Result files are available at http://wilkinsonlab.info/SuppMat/GarciaJimenez/ISMB2016/

## References

Aitchison, J. (1982). The Statistical Analysis of Compositional Data. Journal of the Royal Statistical Society. Series B (Methodological), 44(2), 139–177.

Arumugam, M., Raes, J., Pelletier, E., et al. (2011a). En-terotypes of the human gut microbiome. Nature, 473(7346), 174–80.

Arumugam, M., Raes, J., and al., E. (2011b). Enterotyping tutorial. http://enterotype.embl.de.

Ballou, A. L., Ali, R. A., Mendoza, M. A., Ellis, J. C., Hassan, H. M., Croom, W. J., and Koci, M. D. (2016). Development of the chick microbiome: How early exposure influences future microbial diversity. Frontiers in Veterinary Science, 3(2).

Bellman, R. (1957). A markovian decision process. Indiana Univ. Math. J., 6, 679–684.

Boyd, J. H., Russell, J. A., and Fjell, C. D. (2014). The Meta-Genome of Sepsis: Host Genetics, Pathogens and the Acute Immune Response. Journal of Innate Immunity, 6(3), 272–83.

Brotman, R. M., Ravel, J., Cone, R. A., and Zenilman, J. M. (2010). Rapid fluctuation of the vaginal microbiota measured by Gram stain analysis. Sexually transmitted infections, 86(4), 297–302.

Brotman, R. M., Shardell, M. D., Gajer, P., Tracy, J. K., Zenilman, J. M., Ravel, J., and Gravitt, P. E. (2014). Interplay between the temporal dynamics of the vaginal microbiota and human papillomavirus detection. The Journal of infectious diseases, 210(11), 1723–1733.

Capan, M., Ivy, J. S., Wilson, J. R., and Huddleston, J. M. (2015). A stochastic model of acute-care decisions based on patient and provider heterogeneity. Health care management science.

Chads, I., Chapron, G., Cros, M.-J., Garcia, F., and Sab-badin, R. (2014). MDPtoolbox: a multi-platform toolbox to solve stochastic dynamic programming problems. Ecography, 37(9), 916–920.

Costea, P. I., Zeller, G., Sunagawa, S., and Bork, P. (2014). A fair comparison. Nat Meth, 11(4), 359.

David, L. A. (2014). Normalizing microbiota time-series data. http://nbviewer.ipython.org/github/ladavid/mittimeseries/blob/master/NormalizeDemo.ipynb.

David, L. A., Materna, A. C., Friedman, J., Campos-Baptista, M. I., Blackburn, M. C., Perrotta, A., Erdman, S. E., and Alm, E. J. (2014). Host lifestyle affects human microbiota on daily timescales. Genome biology, 15(7), R89.

Ding, T. and Schloss, P. D. (2014). Dynamics and associations of microbial community types across the human body. Nature, 509(7500), 357–60.

Dominguez-Bello, M. G., Blaser, M. J., Ley, R. E., and Knight, R. (2011). Development of the human gastrointestinal microbiota and insights from high-throughput sequencing. Gastroenterology, 140(6), 1713–1719. Inflammatory Bowel Disease: An Update on Fundamental Biology and Clinical Management.

Faust, K., Lahti, L., Gonze, D., de Vos, W. M., and Raes, J. (2015). Metagenomics meets time series analysis: unraveling microbial community dynamics. Current Opinion in Microbiology, 25, 56–66.

Fritz, J. V., Desai, M. S., Shah, P., Schneider, J. G., and Wilmes, P. (2013). From meta-omics to causality: experimental models for human microbiome research. Microbiome, 1(1), 14.

Gajer, P., Brotman, R. M., Bai, G., Sakamoto, J., Schütte, U. M. E., Zhong, X., Koenig, S. S. K., Fu, L., Ma, Z. S., Zhou, X., Abdo, Z., Forney, L. J., and Ravel, J. (2012). Temporal dynamics of the human vaginal microbiota. Science translational medicine, 4(132), 132ra52.

Hekstra, D. R. and Leibler, S. (2012). Contingency and statistical laws in replicate microbial closed ecosystems. Cell, 149(5), 1164–73.

Howard, R. A. (1960). Dynamic Programming and Markov Processes. MIT Press.

Kaufman, L. and Rousseeuw, P. (1990). Finding Groups in Data: An Introduction to Cluster Analysis. Wiley-Interscience.

Larsen, P. E. and Dai, Y. (2015). Metabolome of human gut microbiome is predictive of host dysbiosis. GigaScience, 4(1), 42.

Maechler, M., Rousseeuw, P., Struyf, A., Hubert, M., and Hornik, K. (2015). cluster: Cluster Analysis Basics and Extensions.

McDonald, D., Clemente, J. C., Kuczynski, J., Rideout, J. R., Stombaugh, J., Wendel, D., Wilke, A., Huse, S., Hufnagle, J., Meyer, F., Knight, R., and Caporaso, J. G. (2012). The Biological Observation Matrix (BIOM) format or: how I learned to stop worrying and love the ome-ome. GigaScience, 1(1), 7.

McMurdie, P. J. and Holmes, S. (2013). phyloseq: An R package for reproducible interactive analysis and graphics of microbiome census data. PLoS ONE, 8(4), e61217.

Nugent, R. P., Krohn, M. a., and Hillier, S. L. (1991). Reliability of diagnosing bacterial vaginosis is improved by a standardized method of gram stain interpretation. Journal of Clinical Microbiology, 29(2), 297–301.

Oakley, B. B., Buhr, R. J., Ritz, C. W., Kiepper, B. H., Berrang, M. E., Seal, B. S., and Cox, N. A. (2014). Successional changes in the chicken cecal microbiome during 42days of growth are independent of organic acid feed additives. BMC Veterinary Research, 10(1), 1–8.

Puterman, M. L. (1994). Markov Decision Processes: discrete stochastic dynamic programming. John Wiley and Sons, Inc.

Rousseeuw, P. J. (1987). Silhouettes: A graphical aid to the interpretation and validation of cluster analysis. Journal of Computational and Applied Mathematics, 20, 53–65.

Sonnenberg, F. A. and Beck, J. R. (1993). Markov models in medical decision making: a practical guide. Medical decision making: an international journal of the Society for Medical Decision Making, 13(4), 322–38.

Voigt, A. Y., Costea, P. I., Kultima, J. R., Li, S. S., Zeller, G., Sunagawa, S., and Bork, P. (2015). Temporal and technical variability of human gut metagenomes. Genome Biology, 16(1), 73.

